# EasyPubPlot: a shiny web application for rapid omics data exploration and visualization

**DOI:** 10.1101/2024.11.26.625339

**Authors:** Nguyen Tran Nam Tien, Nguyen Quang Thu, Dong Hyun Kim, Seongoh Park, Nguyen Phuoc Long

## Abstract

Computational toolkits for data exploration and visualization from widely used omics platforms often lack flexibility and customization. While many tools generate standardized output, advanced programming skills are necessary to create high-quality visualizations. Therefore, user-friendly tools that simplify this crucial yet time-consuming step are essential. We developed EasyPubPlot (Easy Publishable Plotting), a straightforward, easy-to-use, user experience-oriented, open-source, and shiny web application along with its associated R package to streamline data exploration and visualization for functional omics-empowered research. EasyPubPlot generates publishable scores plots, volcano plots, heatmaps, box plots, dot plots, and bubble plots with minimal necessary steps. The tool was designed to guide new users for accurate and efficient navigation. Step-by-step tutorials for each type of plot are also provided. Herein, we demonstrated EasyPubPlot’s competent functionality and versatility by showcasing metabolomics and transcriptomics data. Collectively, EasyPubPlot reduces the gap between data analysis and stunning visualization, thereby diminishing friction and focusing on science. The app can be downloaded and installed locally (https://github.com/Pharmaco-OmicsLab/EasyPubPlot) or used through a web application (https://pharmaco-omicslab.shinyapps.io/EasyPubPlot).

## 1. INTRODUCTION

User-friendly, easy-to-use, and widely supportive applications have become of unparalleled importance for functional omics data exploration and analysis ^1,2^. They have been continuously developed, improved, and updated ^3,4^. These platforms streamline typical analytical workflows, from data pre-processing and exploration to functional and pathway analyses. Besides analytical modules, these tools provide valuable visualizations throughout the analysis pipeline. However, by default, the resulting plots of the statistical analyses often lack flexibility, and the customization is limited, making them less suitable for scientific publication. Although the standardized output from the data analysis tools can be used to generate scientific plots to meet the user’s expectations, this often requires a high level of visualization and programming expertise. Therefore, user-friendly tools that make this crucial but challenging, time-consuming, and high-impact step are essential. Recent efforts have been made to develop and implement these on-demand applications. Despite their potential, these tools are frequently hindered by steep learning curves, limited customization options, and a lack of intuitive interfaces, restricting their adoption by a broader range of researchers ^5–11^. There is an urgent need to develop a highly customizable and user-friendly tool to address this challenging issue.

From our validated in-house visualization strategies, we developed a shiny web application and a companion R package (EasyPubPlot – Easy Publishable Plotting) for the most commonly used and expected plots from recommended analyses for a typical exploratory and validatory functional omics study. EasyPubPlot provides the scores plots, volcano plots, heatmaps, and box plots. Furthermore, dot and bubble plots are also included, which are necessary to effectively present data needed for functional interpretation. Therefore, transcriptomics, proteomics, and metabolomics studies are highly beneficial from the tool. We designed the tool with user-oriented experiences with buttons to guide new users for accurate and efficient navigation. Step-by-step tutorials were also provided. Input data for EasyPubPlot were standard output data from widely used omics applications such as MetaboAnalyst (**Figure 1**). EasyPubPlot is expected to bridge the gap between data analysis and impactful visualization. The uniqueness and versatility of EasyPubPlot are highlighted when placed in perspective with the current scientific app ecosystem (**Table 1**). After a few clicks, users can generate high-quality plots tailored to their specific needs. We focused on metabolomics data to showcase EasyPubPlot’s ability to streamline the visualization process for various omics studies.

**Figure 1.**
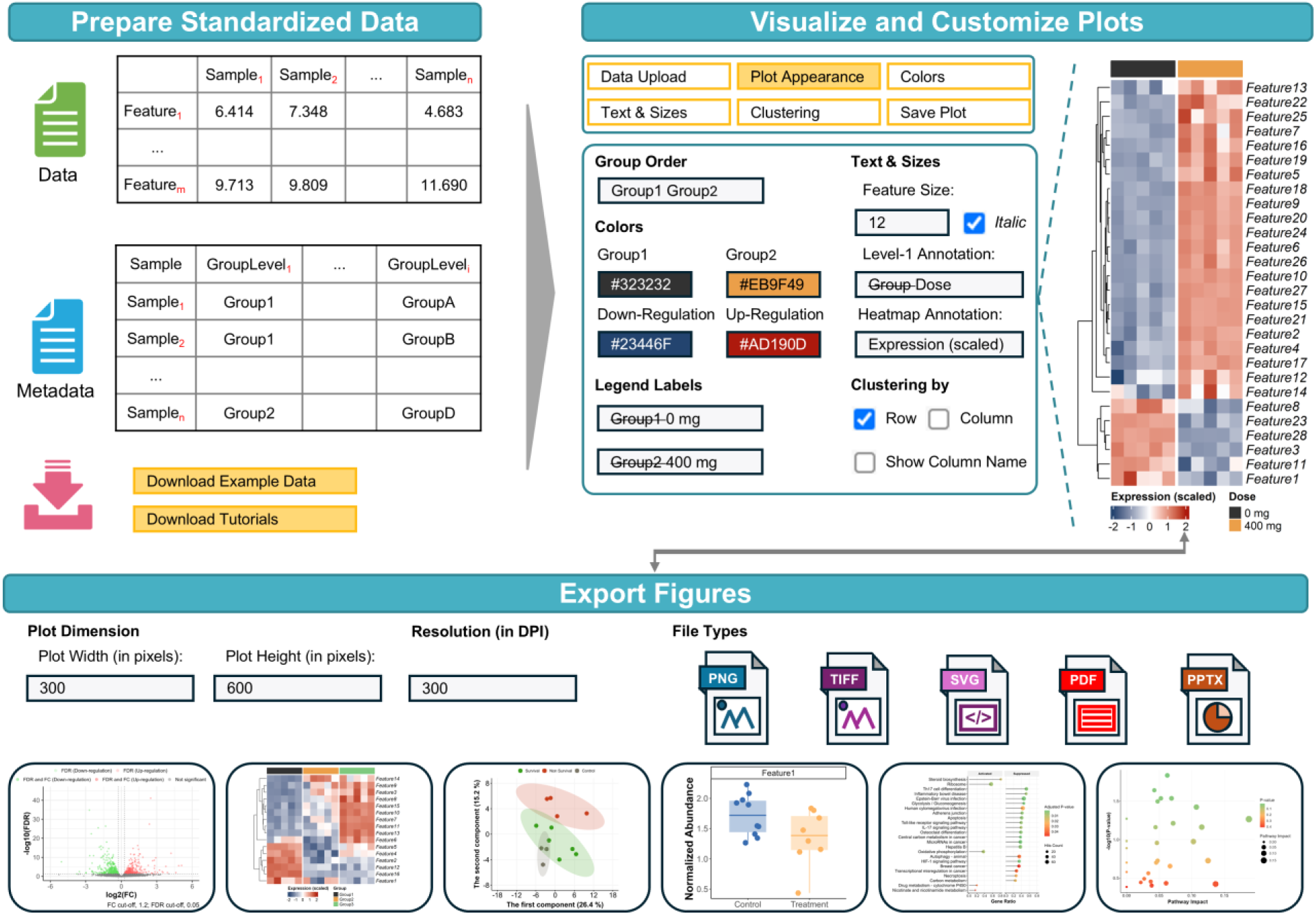
The main functionality of EasyPubPlot with a heatmap plot example. A total of six modules were developed in the first release. EasyPubPlot uses input data as standard output from widely used platforms such as MetaboAnalyst, thereby reducing efforts for data formatting. After uploading data, publishable and ready-to-use plots are shown. Users can easily customize the plot for further use if needed. The plots shown in the user interfaces are identical to downloaded plots, making them easy to customize.

**Table 1.** Comparison with other tools.

|  | ClustVis | VolcaNoseR | GraphBio | Visual Omics | ImageGP 2 | Evenn | EasyPubPlot |
| --- | --- | --- | --- | --- | --- | --- | --- |
| Use generic input | ✓ | ✓ | ✓ | ✓ | ✓ | ✓ | ✓ |
| Provide detailed tutorials | ✓ |  |  | ✓ | ✓ |  | ✓ |
| Support multiple plots in typical explanatory omics studies |  |  | ✓ | ✓ | ✓ |  | ✓ |
| Generate publishable plots | ✓ | ✓ | ✓ | ✓ | ✓ | ✓ | ✓ |
| Customize multiple parameters simultaneously | ✓ | ✓ |  | ✓ | ✓ | ✓ | ✓ |
| Customize multiple parameters interactively | ✓ | ✓ |  |  |  | ✓ | ✓ |
| Export multiple file formats | ✓ | ✓ | ✓ | ✓ | ✓ | ✓ | ✓ |
| Provide online on-demand service | ✓ | ✓ | ✓ | ✓ | ✓ | ✓ | ✓ |
| Support local installation | ✓ | ✓ | ✓ |  | ✓ |  | ✓ |

## 2. IMPLEMENTATION

### 2.1. How to run EasyPubPlot

**Table 2** shows the code metadata. EasyPubPlot is released as an online web application at https://pharmaco-omicslab.shinyapps.io/EasyPubPlot. The open-source code is readily available at https://github.com/Pharmaco-OmicsLab/EasyPubPlot. We also release EasyPubPlot to an R package and distribute it to GitHub to run the app locally. We encourage users to report any issues through GitHub (https://github.com/Pharmaco-OmicsLab/EasyPubPlot/issues).

**Table 2.** Code metadata of EasyPubPlot.

|  |  |
| --- | --- |
| Link to developer manual | https://github.com/Pharmaco-OmicsLab/EasyPubPlot |
| Legal code license | MIT |
| Code versioning system used | git |
| Software code languages, tools, and services used | R <sup>15</sup>, R-Shiny <sup>16</sup>, and https://shinyapps.io/ |
| Compilation requirements, operating environments, and dependencies | R no compilation needed; Dependencies: tidyverse <sup>17</sup>, ggthemes <sup>18</sup>, grid <sup>15</sup>, scales <sup>19</sup>, shiny <sup>16</sup>, shinyWidgets <sup>20</sup>, colourpicker <sup>21</sup>, bslib <sup>22</sup>, shinytoastr<sup>23</sup>, BiocManager <sup>24</sup>, circlize <sup>25</sup>, EnhancedVolcano <sup>26</sup>, and ComplexHeatmap <sup>27</sup> |

When starting, EasyPubPlot has user-oriented buttons to navigate to target panels (**Figure 2A-B**). Additionally, a step-by-step tutorial is provided (**Figure 2B**). This helps avoid confusion for new users during their first trial.

**Figure 2.**
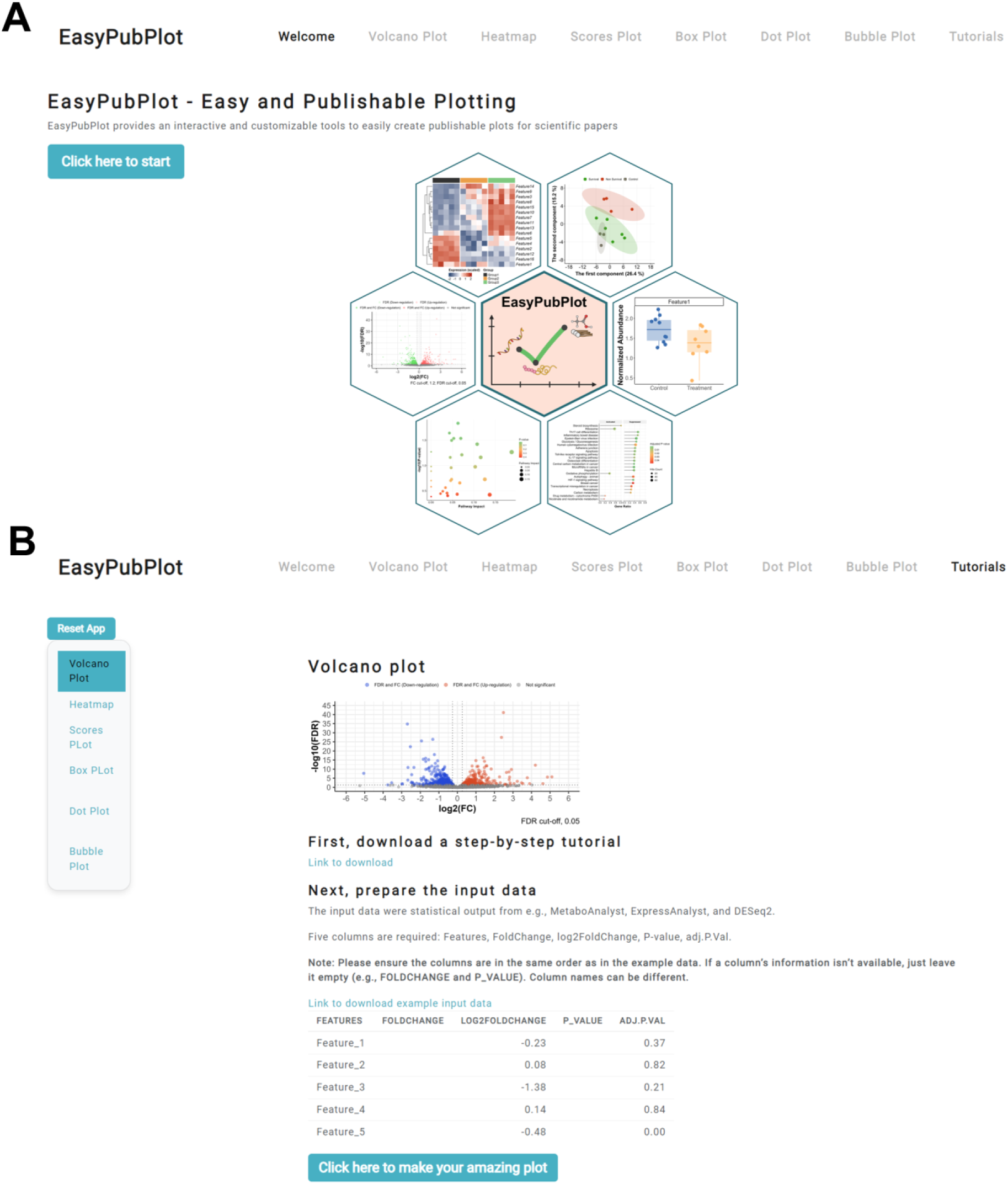
EasyPubPlot view with user experience-oriented buttons and step-by-step tutorials. (A) Welcome page. (B) Tutorials page for volcano plot.

### Inputs and outputs

EasyPubPlot uses standard outputs from widely used applications, such as MetaboAnalyst and ExpressAnalyst ^3,4^. Therefore, minor formatting for input data is required. Users can download example input data directly from EasyPubPlot app.

Regarding the outputs, all plots can be exported with customizable width, height, resolution, and format. When users change the width and height in the user interface, the plots change accordingly, and the downloaded plots have identical visualizations (**Figure 1**). EasyPubPlot provides four formats, including .png, .tif, .svg, and .pdf. The .svg format can be used with other tools for additional customization.

## 3. FUNCTIONALITY AND SHOWCASE

We used an example metabolomics data from MetaboAnalyst and performed a typical analysis workflow using MetaboAnalyst (https://www.metaboanalyst.ca/MetaboAnalyst/) ^3^. For scores plots, volcano plots, heatmap, and box plots, the LC-MS peak intensity table (https://api2.xialab.ca/api/download/metaboanalyst/lcms_table.csv) for mice spinal cord samples with wild-type and knock-out groups, was used (Saghatelian *et al.* ^12^). Data underwent median normalization, log transformation, and Pareto scaling in cases of scores plot visualization. For volcano plots, heatmap, and box plots, data were normalized by median and log-transformed. An example metabolites list from MetaboAnalyst was used for bubble plots and dot plots (https://www.metaboanalyst.ca/MetaboAnalyst/upload/PathUploadView.xhtml). A pathway analysis module with Fisher’s exact test and out-degree centrality was utilized to showcase bubble plot and dot plot. Output from data analysis was used as input for EasyPubPlot.

In the last section, we also demonstrated the versatility of EasyPubPlot by using transcriptomics example data from ExpressAnalyst (https://www.expressanalyst.ca/ExpressAnalyst/resources/data/test/bmd_BRBZ_2w_entrez.txt).

The data were normalized and log-transformed microarray data of rat liver after two weeks of bromobenzene exposure with doses of 0 (control), 25, 100, 200, 300, and 400 mg/kg per day – Dean *et al.* ^13^. For principal component analysis (PCA), data were further scaled by the Pareto method on MetaboAnalyst. ExpressAnalyst was used for statistical analysis and functional analysis. The limma-based method was used for the pairwise analysis between the 400 mg/kg/day group and the control group. The gene set enrichment analysis (GSEA) method with a Kyoto Encyclopedia of Genes and Genomes (KEGG) knowledgebase was used with a fold change-based pre-rank gene list. Output from the data analysis was used as input for EasyPubPlot.

It should be noted that the case studies, comparison, and selected features for heatmap and box plots, were chosen only for visualization purposes.

### 3.1. Scores plot

**Figure 3A-B** shows the scores plot (for principal component analysis (PCA) and partial least squares-discriminant analysis (PLS-DA)) drawn by EasyPubPlot. The scores plot modules also require metadata. The metadata is straightforward to prepare and is reusable for other modules (i.e., heatmap and box plot).

**Figure 3.**
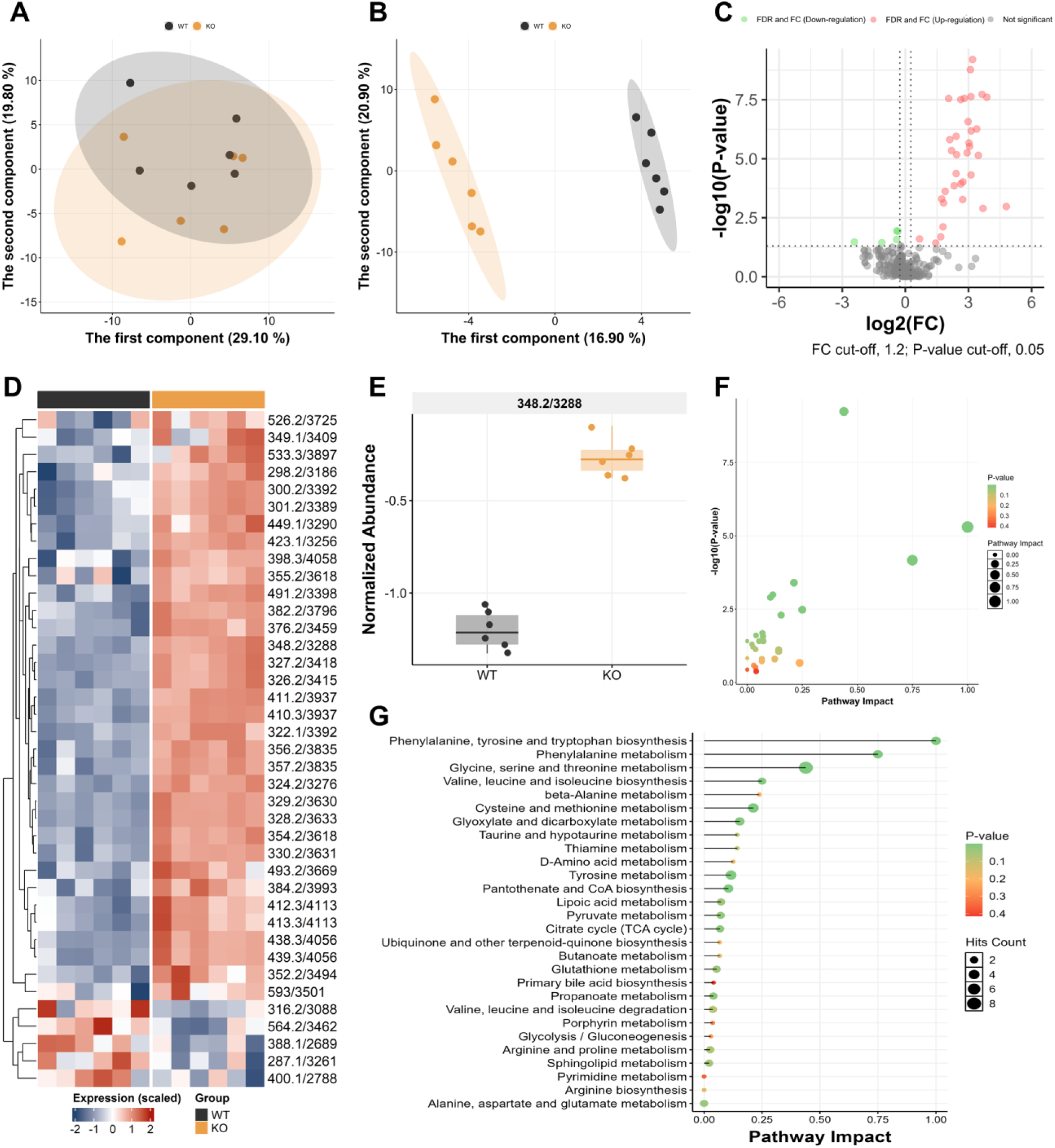
EasyPubPlot showcase for metabolomics. (A) PCA scores plot. (B) PLS-DA scores plot. (C) Volcano plot. (D) Heatmap. (E) Box plot. (F) Bubble plot. (G) Dot plot. Example data were selected only for visualization purposes.

### 3.2. Volcano plot

Next, a volcano analysis was performed to obtain the output for EasyPubPlot. Users need to prepare the input data containing (adjusted) P-value and (log2) fold change of all features in the data. In MetaboAnalyst, this could be done by setting no threshold, i.e., set fold change threshold equal to 1 and a very high (adjusted) P-value cut-off (e.g., 1000) (**Figure 3C**). The P-value or adjusted P-value (highly recommended for exploratory data analysis) could be used for the volcano plot (**Figure 3C**).

### 3.3. Heatmap

Top significant or targeted annotated features could be visualized throughout the heatmap module (**Figure 3D**). The heatmap module requires two input data, i.e., intensity data and metadata. Since there are no pre-processed functions in EasyPubPlot, the intensity data needed to be treated in advance, e.g., via median normalization and log transformation. For example, the data.normalized.csv file exported from the data analysis was used as input for EasyPubPlot (**Figure 3D**).

### 3.4. Box plot

The box plot module (**Figure 3E**) also allows visualization of highly ranked significant or targeted features. The input data (intensity and metadata) are identical to those in the heatmap module. Therefore, the box plot module allows us to visualize the relative abundance of multiple metabolites simultaneously.

### 3.5. Bubble plot

We used the example metabolites list from the pathway analysis module of MetaboAnalyst as an example data to showcase for bubble plot and dot plot (https://www.metaboanalyst.ca/MetaboAnalyst/upload/PathUploadView.xhtml). The output was pathway_results.csv and could be used directly in the bubble plot module for visualization (**Figure 3F**). By default, this module used metabolomics data and P-value for the visualization.

### 3.6. Dot plot

The results from pathway analysis could also be visualized by a dot plot (**Figure 3G**), inspired by the dot plot from the clusterProfiler R package ^14^.

### 3.7. Showcases for transcriptomics data

The case study for using EasyPubPlot for transcriptomics data is illustrated in **Figure 4** for 5 modules, including PCA scores plot (**Figure 4A**), volcano plot (**Figure 4B**), heatmap (**Figure 4C**), boxplot (**Figure 4D**), and dot plot (**Figure 4E**).

**Figure 4.**
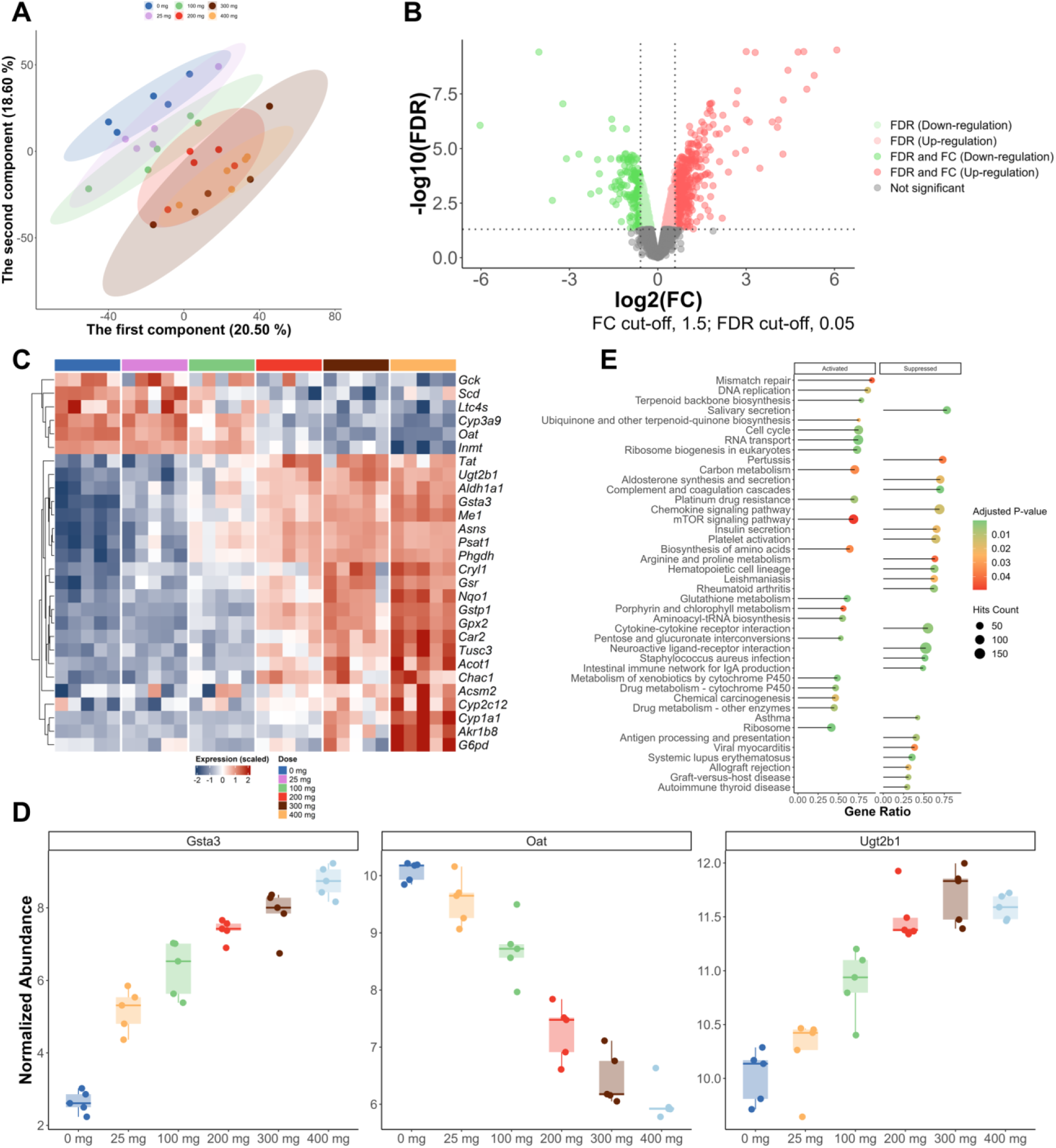
EasyPubPlot showcases for transcriptomics. (A) PCA scores plot. (B) Volcano plot. (C) Heatmap. (D) Box plot. (E) Dot plot. Example data were selected only for visualization purposes.

## 4. CONCLUSIONS

EasyPubPlot is open-source omics software that offers a user-friendly platform for generating high-quality figures with minimal effort. Crucial visualization modules are provided within a single, intuitive, user experience-oriented interface, saving time and reducing the effort required to generate publishable, high-quality figures. The app accelerates data exploration and visualization, enabling researchers to communicate their findings effectively. Furthermore, EasyPubPlot’s modular design and open-source nature facilitate continuous improvement. Future developments may include 3D visualization, interactive plots, and support for emerging omics technologies.

## Data Availability Statement

The metabolomics data originally reported by Saghatelian *et al.* ^12^ and available for download from MetaboAnalyst (https://api2.xialab.ca/api/download/metaboanalyst/lcms_table.csv).

The transcriptomics data originally reported by Dean *et al.* ^13^ and available for download from ExpressAnalyst (https://www.expressanalyst.ca/ExpressAnalyst/resources/data/test/bmd_BRBZ_2w_entrez.txt).

Example input data are available in the app – freely available at https://pharmaco-omicslab.shinyapps.io/EasyPubPlot and at https://github.com/Pharmaco-OmicsLab/EasyPubPlot.

## Author Contributions

N.P.L. conceptualized the study. N.T.N.T. developed and validated the software. N.Q.T. and N.T.N.T. prepared the tutorials and case studies. D.H.K., S.P., N.P.L. supervised the project. N.T.N.T. and N.P.L. drafted the first version of the manuscript. N.P.L., S.P., D.H.K., and N.Q.T. reviewed and edited the manuscript. All authors approved the final version of the manuscript.

## ACKNOWLEDGMENTS

We thank Nguyen Ky Phat and Nguyen Thi Hai Yen for in-house code development and app testing.

